# The genetic basis of cis-regulatory divergence between the subspecies of cultivated rice (*Oryza sativa*)

**DOI:** 10.1101/511550

**Authors:** Malachy T Campbell, Qian Du, Kan Liu, Sandeep Sharma, Chi Zhang, Harkamal Walia

**Affiliations:** Department of Agronomy and Horticulture, University of Nebraska Lincoln, Lincoln, NE, USA 68583; School of Biological Sciences, University of Nebraska Lincoln, Lincoln, NE, USA 68583; Marine Biotechnology and Ecology Division, CSIR-CSMCRI, Bhavnagar, Gujarat, India

**Keywords:** Transcriptome, rice, expression QTL, genome-wide association, genomics

## Abstract

Cultivated rice consists of two subspecies, *Indica* and *Japonica*, that exhibit well-characterized differences at the morphological and genetic levels. However, the differences between these subspecies at the transcriptome level remains largely unexamined. Here, we provide a comprehensive characterization of transcriptome divergence and cis-regulatory variation within rice using transcriptome data from 91 accessions from a rice diversity panel (RDP1). The transcriptomes of the two subspecies of rice are highly divergent. The expression and genetic diversity was significantly lower within *Japonica* relative to *Indica*, which is consistent with the known population bottleneck during *Japonica* domestication. Moreover, 1,860 and 1,325 genes showed differences in heritability in the broad and narrow sense respectively, between the subspecies, which was driven largely by environmental and genetic effects rather than differences in phenotypic variability. We leveraged high-density genotypic data and transcript levels to identify cis-regulatory variants that may explain the genetic divergence between the subspecies. We identified significantly more eQTL that were specific to the *Indica* subspecies compared to *Japonica*, suggesting that the observed differences in expression and genetic variability also extends to cis-regulatory variation. We next explored the potential causes of this cis-regulatory divergence by assessing local genetic diversity for cis-eQTL. Local genetic diversity around subspecies-specific cis-eQTL was significantly lower than genome-wide averages in subspecies lacking the eQTL, suggesting that selective pressures may have shaped regulatory variation in each subspecies. This study provides the first comprehensive characterization of transcriptional and cis-regulatory variation in cultivated rice, and could be an important resource for future studies.

## Introduction

Cultivated rice consists of two subspecies: *Indica* and *Japonica*. *Indica* varieties are cultivated throughout the tropics, and account for the majority of rice production worldwide. *Japonica* varieties, on the other hand, are grown in both tropical and temperate environments, and only account for approximately 20% of rice production.

Although the domestication history of rice remains a contested topic, the most current research collectively suggests that rice was domesticated at least twice from two geographically and ecologically distinct subpopulations of *Oryza rufipogon*. The unique environmental pressures in these distinct regions, as well as preferences by early farmers for grain characteristics has resulted in large morphological and physiological differences between the two subspecies. These differences have been recognized for centuries, as evidenced by references of Keng and Hsein types of rice found in records from the Han Dynasty in China (Oka et al., 1991).

The unique natural and agronomic selection pressures placed on the wild progenitors and early proto-domesticates resulted in drastic changes at the genetic level. Work by Huang et al. (2012b) showed considerable reduction in genetic diversity in *Indica* and *Japonica* compared with *O. rufipogon*. Such drastic reductions in genetic diversity are common following domestication. Moreover, the transition from an out-crossing/heterogamous nature of *O. rufipogon* to the autogamous breeding system of cultivated rice likely led to greater partitioning of genetic diversity among the two subspecies, and further differentiation of the two groups. These large genetic differences have been recognized for nearly a century as hybrids between *Indica* and *Japonica* exhibit low fertility (Kato, 1928). More recently, these genetic differences have been realized with the availability of high density molecular markers and full genome sequences for both *Indica* and *Japonica* (Ding et al., 2007; Goff et al., 2002; Yu et al., 2002; Feltus et al., 2004; Stein et al., 2018; Koide et al., 2018; Schatz et al., 2014; Huang et al., 2008; Wang et al., 2014; Huang et al., 2012b). For instance Ding et al. (2007) showed that approximately 10% of the genes in the *Indica* and *Japonica* genomes showed evidence of presence-absence variation or asymmetrical genomic locations. Several other studies have highlighted genetic differences between the subspecies as structural variants differences, gene acquisition and loss, transposable element insertion and single nucleotide polymorphisms (Goff et al., 2002; Yu et al., 2002; Feltus et al., 2004; Stein et al., 2018; Koide et al., 2018; Schatz et al., 2014; Huang et al., 2008; Wang et al., 2014; Huang et al., 2012b).

While the morphological and genetic differences of *Indica* and *Japonica* have received considerable attention, few studies have investigated the divergence between the two subspecies at transcriptome level (Walia et al., 2007; Lu et al., 2010; Jung et al., 2013). Walia et al. (2007) utilized genome-wide expression profiling to characterize the transcriptional responses for two *Indica* and *Japonica* cultivars to salinity. This study was performed to elucidate the mechanisms underlying the contrasting responses to stress exhibited by the cultivars, rather than examine the transcriptional difference between the subspecies. Moreover, separating genotypic differences from subspecies differences is not feasible with the low number of cultivars used in these studies. Lu et al. (2010) compared transcriptional profiles of two *Indica* accessions and a single *Japonica* accessions and identified many novel transcribed regions, highlighted alternative splicing differences, and differentially expressed genes between accessions. Although these studies provided insights into the transcriptional differences between *Indica* and *Japonica*, given the small sample size of the study it has limited scope for extending conclusions to a population level. Jung et al. (2013) leveraged the large number of public microarray databases to compare transcriptional diversity between the two subspecies. The 983 publicly available Affymetrix microarrays were classified into *Indica* and *Japonica* subspecies based on the cultivar name. This study showed that considerable differences in expression levels were evident between the two subspecies. However, considerable information is likely lost due to the heterogeneity in sample types (e.g. tissue, developmental stage) and varying growth conditions. Thus, a more highly controlled study that utilized a larger panel with genotypic information would provide greater insight into the differences in expression levels, as well as provide a mechanism for connecting transcriptional differences between the two subspecies with genetic variation.

The objective of this study is to examine genetic basis of the transcriptional variation at a population level within the *O. sativa* species. By combining population and quantitative genetics approaches, we aim to elucidate the genetic basis of transcriptional divergence between the two subspecies. To this end, we generated transcriptome data using RNA sequencing on shoot tissue for a panel of 91 diverse rice accession selected from the Rice Diversity Panel1 (RDP1) (Zhao et al., 2011; Famoso et al., 2011; Eizenga et al., 2014). Here, we show that transcriptional diversity between *Indica* and *Japonica* subspecies is consistent with diversity at the genetic level. Moreover, we connect transcriptional differences between the two subspecies with divergent patterns of *cis*-regulatory variation and show that the absence of many *cis*-regulatory variants are due to unique selective pressures experienced by each subspecies. This study is the first to document the transcriptional divergence between the major subspecies of cultivated rice at a population level, and provides insight into the genetic mechanisms that have shaped this transcriptional divergence.

## Materials and Methods

### Plant materials and growth conditions

This study used 91 diverse accessions from the Rice Diversity Panel1 (RDP1) (Famoso et al., 2011; Zhao et al., 2011; Eizenga et al., 2014). Seeds were obtained from the USDA-ARS Dale Bumpers Rice Research Center. The 91 accessions consisted of 13 *admixed*, 2 *aromatic*, 9 *aus*, 23 *indica*, 21 *temperate japonica*, and 23 *tropical japonica* accessions.

Seeds were dehusked manually and germinated in the dark for two days at 28°C in a growth cabinet (Percival Scientific), and were exposed to light (120 *µ*mol *m*^-2^*s*^-1^) twelve hours before transplanting to acclimate them to the conditions in the growth chamber. The seeds were transplanted to 3.25” × 3.25” × 5” pots filled with Turface MVP (Profile Products) in a walk-in controlled environment growth chamber (Conviron). The pots were placed in 36” × 24” × 8” tubs, that were filled with tap water. Fours days after transplanting the tap water was replaced with half-strength Yoshida solution (Yoshida et al., 1976) (pH 5.8). The pH of the solution was monitored twice daily and was recirculated from a reservoir beneath the tubs to the growth tubs. The temperatures were maintained at 28°C and 25°C in day and night respectively and 60% relative humidity. Lighting was maintained at 800 *µ*mol *m*^-2^*s*^-1^ using high-pressure sodium lights (Phillips).

### RNA extraction and sequencing

Ten days after transplant, aerial parts of the seedlings were excised from the roots and frozen immediately in liquid nitrogen. The samples were ground with Tissuelyser II (Invitrogen) and total RNA was isolated with RNAeasy isolation kit (Qiagen) according to manufacturer’s instructions. On-column DNAse treatment was performed to remove genomic DNA contamination (Qiagen). Sequencing was performed using Illumina HiSeq 2500. Sixteen RNA samples were combined in each lane. Two biological replicates were used for each accession.

### Sequence alignment, expression quantification, and differential expression analysis

Quality control for raw reads was performed using the package FastQC (Andrews et al., 2010). The Illumina 101-bp single-end reads were screened and trimmed using Trimmomatic to ensure each read has average quality score larger than 30 and longer than 15 bp, and were aligned to the rice genome (Oryza sativa MSU Release 6.0) using TopHat (v.2.0.10), allowing up to two base mismatches per read. Reads mapped to multiple locations were discarded (Trapnell et al., 2009; Bolger et al., 2014). The number of reads for each gene sequence was counted using the HTSeq-count tool with the “union” resolution mode (Anders et al., 2015). For down-stream genetic analyses, a variance stabilized transformation was performed on raw read counts to provide approximately homoskedastic values in DEseq2 (Love et al., 2014).

To identify genes that exhibited differential expression between the two subspecies, a mixed linear model was fit that included subspecies as the main fixed effect and accession as a random effect in lme4 (Bates et al., 2015). This ‘full’ model was compared to a redcued model the lacked subspecies as a fixed effect using a likelihood-ratio test. Prior to differential expression analysis, expression levels were quantile normalized to ensure a Gaussian distribution. Benjamini and Hochberg’s method was used to control the false discovery rate, and genes with an FDR *≤* 0.001 were considered differentially expressed (Benjamini and Hochberg, 1995).

Genes showing differences in presence-absence expression variation (PAV) was determined using a mixed-effects logistic regression model. Briefly, for each sample the expressed genes (number of reads ¿ 10) were assigned 1, while those with 10 or less reads were assigned a 0. A logistic regression model was fit using the ‘glmer’ function in ‘lme4’ and included subspecies as a fixed effect and accession as random (Bates et al., 2015). The significance of the fixed effect of subspecies was determined by comparing the full model above with a reduced model that lacked subspecies using a likelihood-ratio test. Benjamini and Hochberg’s method was used to control the false discovery rate, and genes with an FDR *≤* 0.001 were considered as having presence-absence expression variation (Benjamini and Hochberg, 1995).

### Subspecies classification

The 91 accessions were classified into two subspecies using the software STRUCTURE (Pritchard et al., 2000). Briefly, the software was run using the 44k SNP data, assuming two subpopulations (K=2), with 20000 MCMC replicates and a burn-in of 10000 MCMC replicates.

### Expression and genetic diversity analyses

Principle component analysis of gene expression was conducted for the 91 accessions using 22,675 genes after variance stabilizing transformation. For, PCA of SNP data the 44k dataset described by Zhao et al. (2011) was used. SNPs with a MAF *<* 0.10 were removed prior to PCA analysis.

The coefficient of variation (CV) was used to estimate the diversity in gene expression within the *Indica* and *Japonica* subspecies. Prior to estimating CV genes with low expression (i.e. those with read counts of *≤* 10 in *≥* 20% of the samples) were removed, leaving a total of 22,503 genes in *Japonica* and 21,719 genes in *Indica*. For the estimation of *π*, SNPs were extracted for each subspecies and SNPs with MAF ¡ 0.05 were excluded. In total 201,891 SNPs were retained for *Indica* and 161,715 for *Japonica*. *π* was estimated at each SNP using the site-pi function in VCFtools (Danecek et al., 2011).

### Heritability estimates

Heritability, both in the broad (*H*^2^) and narrow sense (*h*^2^), was estimated across subspecies for 22,675 genes that were expressed in both *Indica* and *Japonica*. To estimate *H*^2^ a mixed model was fit using lme4 where accession was considered a random effect, and significance of *H*^2^ was assessed using a restricted likelihood-ratio test in the RLRTsim package (Bates et al., 2015; Scheipl et al., 2008). Benjamini and Hochberg’s method was used to control the false discovery rate, and genes with an FDR *≤* 0.001 were considered to have significant genetic variability (Benjamini and Hochberg, 1995). To assess hertiability in the narrow sense (*h*^2^) a mixed model was fit in asreml-R (Butler et al., 2009). Briefly, a genomic relationship matrix (G) was estimated according to VanRaden (2008) using the approximately 36,901 SNPs described by Zhao et al. (2011). G is estimated as 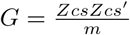, where *Z*_*sc*_ is the centered and scaled marker matrix and *m* is the number of markers. A likelihood-ratio test was used to assess significance and Benjamini and Hochberg’s method was used to control the false discovery rate. Genes with an FDR *≤* 0.001 were considered to have significant genetic variability (Benjamini and Hochberg, 1995).

Heritability was assessed within subspecies using the same approaches as described above. However, due to the unequal sample size for the *Indica* and *Japonica* subspecies, a random set of 35 *Japonica* accessions were selected. Genes showing low expression (¡ 10 reads in ¡ 20% of samples) in either subspecies were removed prior analysis, leaving 22,444 genes in *Japonica* and 22,068 genes in *Indica*.

### Assessing differences in genetic variability between subspecies

To identify genes showing significant differences in genetic variability (*H*^2^ or *h*^2^) between subspecies, a permutation approach was used. Here, the 91 accessions were randomly partitioned into two groups of equal size (35 accessions each). Hertiability was estimated as described above and the difference in heritability between each group was calculated. The resampling approach was repeated 100 times for both *H*^2^ and *h*^2^. This process effectively estimated a null distribution of Δ*H*^2^ and Δ*h*^2^ values. The heritability estimates for each subspecies was used to calculate the differences in *H*^2^ and *h*^2^ between the two subspecies as 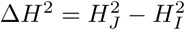 or 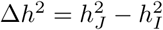. These values were compared with the null distribution to assess significance.

### Joint cis-eQTL analysis

eQTLs were jointly detected using the eQTL-BMA (Bayesian model averaging) described by Flutre et al. (2013) for 26,675 genes and 274,499 SNPs (MAF *>* 0.10) McCouch et al. (2016). Prior to eQTL mapping BLUPs for each gene was calculated and the gene expression level of each gene was transformed into the quantiles of a standard Normal distribution with ties broken randomly. To control for the effects of population structure the first four PCs derived from PCA analysis of 44k SNP dataset were included in the linear model. Briefly, to identify eQTL and control false discovery rate (FDR) a gene-level permutation approach was used within the eQTL-BMA software. Using the eqtlbma bf program, 10,000 permutations were performed with the following settings: –maf 0.1, –nperm 10000, –trick 1, –tricut 10 and –error uvlr. Genes were considered to have an eQTL if the *FDR* < 0.05. These permutations were used to estimate *π*_0_, the probability for a gene to have no eQTL in any subspecies. Here, expression from both *Japonica* and *Indica* samples were analyzed together with the option –error uvlr specified. Next, a hierarchical model with an expectation–maximization algorithm was used to estimate hyper-parameters and configuration probabilities using the eqtlbma hm program. These configurations were *Indica*-specific, *Japonica*-specific, and present in both subspecies. Lastly, the eqtlbma avg bfs program was run to obtain (i) the posterior probability (PP) of a gene to have an eQTL in at least one subspecies, (ii) PP for a SNP to be the causal SNP for the eQTL, (iii) PP for the SNP to be an eQTL, (iv) PP for the eQTL to be present in one subspecies, and (v) PP for the eQTL to be present for a specific configuration. SNP-gene pairs were determined to be specific to a given subspecies or shared if the PP *>* 0.5 for a given configuration.

### Detecting evidence of selection at cis-eQTL

To determine whether the absence of an eQTL was due to of selection, first SNPs from the HDRA dataset within 100kb of each significant eQTL were extracted for the 91 accessions McCouch et al. (2016). For each SNP, nucleotide diversity was determined using the site-pi function in VCFtools and was averaged across the 100kb window (Danecek et al., 2011). Secondly, a genome-wide diversity level was determined for each subspecies. Here, SNPs that were within 100kb of an eQTL were excluded, as well as those that exhibited low diversity in both subspecies (MAF *<* 0.1 in both *Indica* and *Japonica*). Nucleotide diversity was determined as described above for each SNP, and the average was taken for 100kb windows. For each class of eQTL (e.g. *Indica*-specific, *Japonica*-specific, and shared), a two-sided Student’s *t*-test was performed to assess whether the mean *π* was different from the genome-wide average for each subspecies and class of eQTL.

A similar approach was taken for the 3kg data (Alexandrov et al., 2014). For each eQTL SNP, all SNPs within 100kb of the eQTL SNP was extracted from the 4.8M core SNP data. The MAF was determined for each of the 12 subpopulations in the 3kg data, and SNPs that had low diversity (MAF *<* 0.01) in 10 of the 12 subpopulations were excluded from further analyses. As above, *π* was calculated for each site. An average *π* was determined for each subpopulation at each eQTL by taking the average *π* across the 100kb window. To obtain a genome wide average, eQTL regions were excluded and *π* was obtained for each subpopulation by averaging *π* across the 100kb region. Finally, as above a two-sided Student’s *t*-test was performed to assess whether the mean *π* was different from the genome-wide average for each subpopualtion and class of eQTL.

## Results

We selected 91 accessions to represent the genetic diversity within Rice Diversity Panel 1 (RPD1). Using the subpopulation assignment described by Zhao et al. (2011) and Famoso et al. (2011), shoot transcriptome data was generated for 23 *tropical japonica*, 23 *indica*, 21 *temperate japonica*, 13 *admixed*, 9 *aus*, and 2 *aromatic* accessions. Genes with low variance or expression within the expression set were filtered out, as these genes are uninformative for downstream analyses focused on natural variation in gene expression. A total of 25,732 genes were found to be expressed (*>* 10 read counts) in at least one or more of the 91 accessions. This equates to about 46% of the genes present in the rice genome (total of 55,986 genes in MSUv7 build).

### Divergence between the *Indica* and *Japonica* subspecies are evident at the genetic and transcriptional levels

To examine patterns of variation within the transcriptomics data, we performed principle component analysis (PCA) of transcript levels for the 91 accessions. Prior to PCA, lowly expressed genes were removed if they were not expressed (*<* 10 reads) in at least 20% of the samples. This filtering removed approximately 33,311 genes, resulting in a total of 22,675 genes that were used for the principal component analysis based on the normalized read counts. For the genetic analysis, we used 32,849 SNPs. PCA analysis of the expression matrix resulted in a clear separation between the two subspecies along PC1, suggesting a significant transcriptional divergence between *Indica* and *Japonica* (Figure 1C,D). The first PC accounted for approximately 26.8% of the variation in gene expression. While PC1 was able to differentiate between the two subspecies at the transcriptional level, no clear clustering of accessions was observed along other PCs (Figure 1). These results suggest that the the two subspecies of cultivated rice have divergent transcriptomes, but the transcriptomes of the subpopulations are more similar. Consistent with these results observed for PCs 1 and 2, differentiation between the subspecies was clearly evident along PC1 using the genetic (SNP) data alone (Figure 1A,B). However, the clustering of accessions along PCs 2-4 for the SNP data were consistent with those described by Zhao et al. (2011) (Figure 1), and were effective in discerning the two subpopulations in rice. These results collectively suggest that the two subspecies are vastly divergent at genetic and transcriptional level.

**Figure 1.**
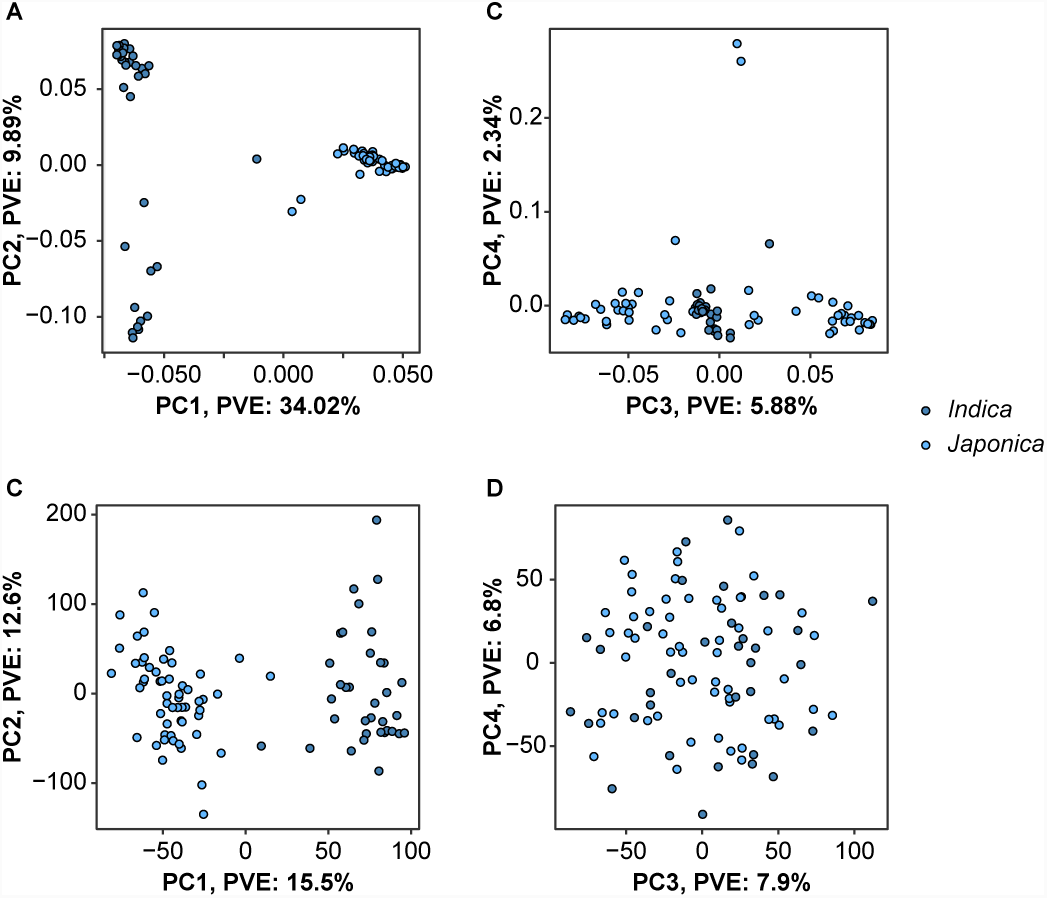
Principle component analysis of markers and gene expression matrices. The top four principle components from PCA analysis of the expression data are pictured in A and B to illustrate the divergence of the major subpopulations in rice. The panels in C and D summarize PCA of genotypic data. PVE: percent variation explained by each component.

### Differential expression analysis reveals contrasting expression between subspecies

To further explore the differences and identify genes that display divergent expression between the two subspecies, the 91 accessions were first classified into *Indica* and *Japonica*-like groups, using the program STRUCTURE with the assumption of two groups and no admixture (Pritchard et al., 2000). A total of 35 accessions were assigned to the *Indica* subspecies, while 56 were assigned to the *Japonica* subspecies. Next, a linear mixed model was fit for each of the 26,675 genes, where subspecies was considered a fixed effect and accession as a random effect. A total of 7,417 genes were found to exhibit contrasting expression between the two subspecies (FDR *≤* 0.001, Supplemental File S1). Of these genes, 4,210 (57%) showed significantly higher expression in *Japonica* relative to *Indica*, while 3,207 (43%) showed higher expression in *Indica* relative to *Japonica*.

This divergent expression levels observed between the two subspecies could be the result of the presence or absence of genes within the subspecies. To this end, we sought to identify genes showing a presence-absence expression variation (PAV). Genes with a read count greater than 10 were considered as expressed and coded as 1 while those with read counts less than 10 were coded as 0. These genes were further filtered, so that genes that were expressed in at least 20%, but no more than 80% of the samples were retained for downstream analyses. A logistic mixed effects model was fit for the 4,263 genes meeting this criteria. In total, 1,980 genes showed evidence of PAV between the two subspecies (*FDR* < 0.001; Supplemental File S1). This analysis, enriched for genes that were expressed at higher frequency in *Japonica* rice compared to *Indica*. For instance, 1,435 genes were found to be expressed at a significantly greater frequency in *Japonica* relative to *Indica*, while only 545 were found to be expressed predominately in *Indica*. Moreover, we detected significant enrichment for GO terms associated stress response GO:00006950)and response to biotic stress (GO:0009607), as well genes with kinase activity (GO:0016301). Within *Indica*-specific genes, only a single GO category was enriched for oxygen binding activity (GO:0019825; Table S1). Moreover, 173 were identified with no evidence of expression in *Indica* while only 18 were identified in *Japonica*. Collectively, these results suggest that the divergence between *Indica* and *Japonica* subspecies may be due, in part, to differences in mean expression levels as well as presence-absence expression variation.

### Japonica subspecies exhibits reduced genetic and transcriptional diversity

Several studies have shown that the unique domestication history of the two subspecies has resulted in large differences in the overall genetic diversity between the two subspecies, with *Indica* being more genetically diverse than *Japonica* Caicedo et al. (2007); Huang et al. (2010, 2012b); Mather et al. (2007). We next explored the variation in gene expression within each subspecies. Two metrics were used to examine the differences in diversity at both the genetic and transcriptional levels within each subspecies: nucleotide diversity (*π*) and the coefficient of variation (CV). Diversity analyses within each subspecies may be influenced by differences in sample size. Since the number of *Japonica* accessions were greater than *Indica*, a subset of 35 *Japonica* accessions were randomly selected for diversity analyses. The results for the full set of 56 *Japonica* accessions are provided as Figure S1.

Expression diversity was estimated using the coefficient of variation (CV) for 22,675 genes. CV was significantly different between the two subspecies (Wilcoxon rank sum test, *p* < 0.0001; Figure 2). The *Indica* subspecies exhibited approximately 12.6% higher expression diversity compared to *Japonica*. On average, CV in the *Indica* subspecies was 3.46, while in the *Japonica* subspecies the mean CV was 3.07. These results suggest that the transcriptional diversity is lower in the *Japonica* subspecies compared to *Indica*. CV estimates using the complete set of *Japonica* accession were similar (CV: 3.46 and 3.10 for *Indica* and *Japonica*, respectively; Figure S1).

**Figure 2.**
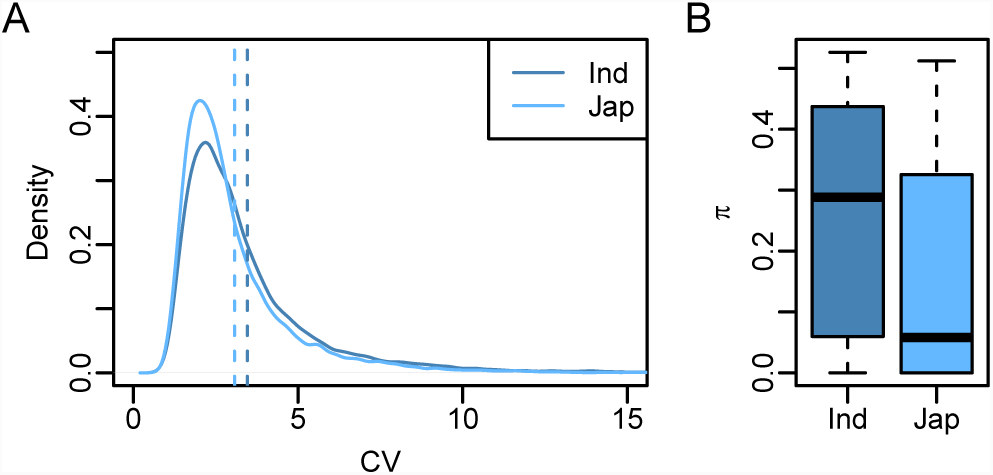
Genetic and expression diversity within *Indica* and *Japonica* accessions. (A) The coefficient of variation was used as an estimate of the diversity in gene expression within each subspecies. A subset of 35 *Japonica* accessions were randomly selected for diversity analyses to ensure that sample sizes were equal between the two subspecies. The vertical dashed lines represent the mean CV within each subspecies. (B) Site-wise nucleotide diversity (*π*) was used as an estimate of the genetic diversity within each of the subspecies using 36,901 SNPs described by Zhao et al. (2011).

Genetic diversity within each subspecies was estimated using *π* for 33,543 SNPs in randomly selected 35 *Indica* and 35 *Japonica* accessions. Similar differences were observed for *π* as CV, however the differences between subspecies was much greater (Wilcoxon rank sum test, *p* < 0.0001; Figure 2). The *Indica* subspecies showed a 64.7% higher nucleotide diversity (*π*) compared to *Japonica*. On average, *π* estimates were 0.26 for *Indica* and 0.17 for *Japonica*. These results are consistent with reports by Huang et al. (2012b) and Garris et al. (2005), and are in agreement with the expression diversity reported above. Together these data suggest that the *Japonica* subspecies exhibits less genetic and transcriptional diversity compared to *Indica*.

### Gene expression is heritable in cultivated rice

The above analyses shows a strong differentiation between the subspecies at transcriptional and genetic levels, and presents a possible linkage between expression and genetic diversity. However, the extent of variation in gene expression that can be accounted by genetic variation is not yet determined. To estimate the extent to which variation in gene expression is under genetic control, a mixed model was fit to the expression of each of the 22,675 genes and the variance between accessions was estimated. The significance of the random *between - accession* term was determined using a likelihood-ratio test. The broad-sense heritability (*H*^2^) was estimated as the proportion of the total variance explained by between-accession variance to total variance. A total of 11,895 genes showed a significant *between - accession* variance (*FDR* < 0.001; *H*^2^ *≥* 0.47), which accounts for approximately 53% of the genes expressed in at least 20% of the samples (Figure 2A; Supplemental File S2). *H*^2^ ranged from 0.97 to 0.47, with 4,606 genes showing highly heritable expression (*H*^2^ *>* 0.75), 7,145 showing moderate *H*^2^ (0.5 *< H*^2^ *≤* 0.75), and the remaining 146 showing low *H*^2^.

To determine the extent to which additive genetic effects could explain variance in gene expression, a genomic relationship matrix was constructed using 32,849 SNPs following VanRaden (2008) and variance components were estimated using a mixed linear model for each gene. A total of 10,125 genes were identified with significant *h*^2^ (Supplemental File S2). Of these, 234 genes had highly heritable expression (*h*^2^ *≥* 0.75), while 2,750 genes showed moderate heritability (0.5 *≤ h*^2^ *<* 0.75) (Figure 3B). An additional 7,141 genes showed low narrow sense heritability (*h*^2^ *<* 0.5). Collectively, these results indicate that a large portion of the rice transcriptome is under genetic control.

**Figure 3.**
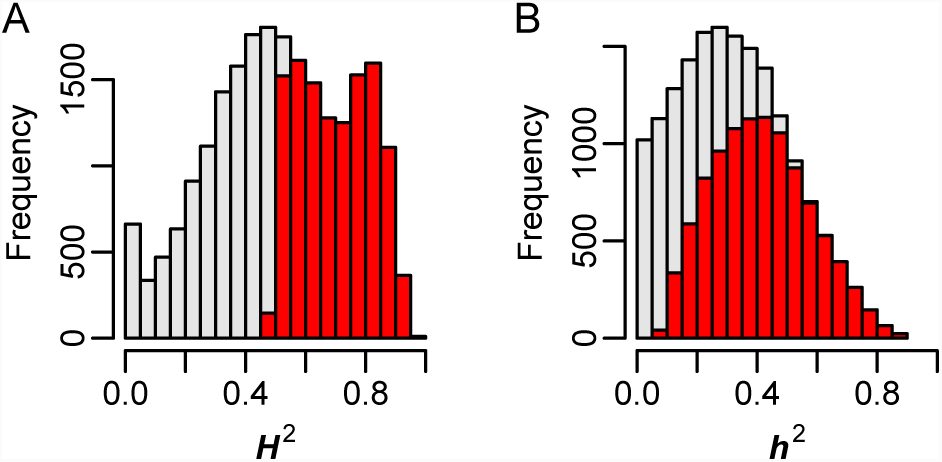
Heritability of gene expression across *O. sativa* subspecies. Distribution of broad-sense heri-tability (*H*^2^) and narrow-sense heritability (*h*^2^) for 22,675 genes are pictured in panels A and B, respectfully. Bars highlighted in red indicate genes with significant genetic effects (*FDR* < 0.001).

### Genetic variability of gene expression is considerably different between subspecies

The analyses above indicate that the two subpopulations differ at the transcriptional and genetic levels, and that for many genes, variation in expression can be explained by genetic effects. We next asked whether the heritability of gene expression is different between the two subspecies. To this end, the expression dataset was partitioned into *Indica* and *Japonica* subsets and genes with low expression in each subspecies were remove (expressed in less than 20% of the samples). Since the number of accessions for the two subspecies are unequal, 35 *Japonica* accessions were randomly sampled to ensure the two samples were of equal size, and the number of genes that were expressed in each subspecies were quantified. Here, a gene was considered expressed if 10 or more reads mapped to the gene in 20% or more of the samples. A total of 22,444 genes were found to be expressed in at least 20% of the samples for the *Japonica* subspecies, while 22,068 were found to be expressed in the *Indica* subspecies. A large number of genes were common to both subspecies (21,166 genes). A total of 1,278 genes were found to be uniquely expressed in *Japonica*, and 902 were found to be uniquely expressed in *Indica*.

A total of 5,005 genes exhibited significant *H*^2^ in *Indica* and 3,338 genes in *Japonica* (*FDR* < 0.001; Supplemental File S3). For these genes, *H*^2^ ranged from 0.67 to 0.98 in *Indica* and 0.67 to 0.97 in *Japonica*. A larger number of genes were identified with significant additive genetic variance, with 6,804 identified in *Indica* and 5,103 found in *Japonica*. For these genes, narrow-sense heritability ranged from 0.201 to 0.953 in *Indica* and 0.220 to 0.948 in *Japonica*. Interestingly, few genes showed significant heritable expression in both subspecies. For instance, only 1,681 and 2,644 genes were found to have significant *H*^2^ and *h*^2^, respectively, in both *Indica* and *Japonica*. Moreover, a comparison of *H*^2^ and *h*^2^ between subspecies showed that for many genes, heritability estimates were considerably different between *Indica* and *Japonica* (Figure 4).

**Figure 4.**
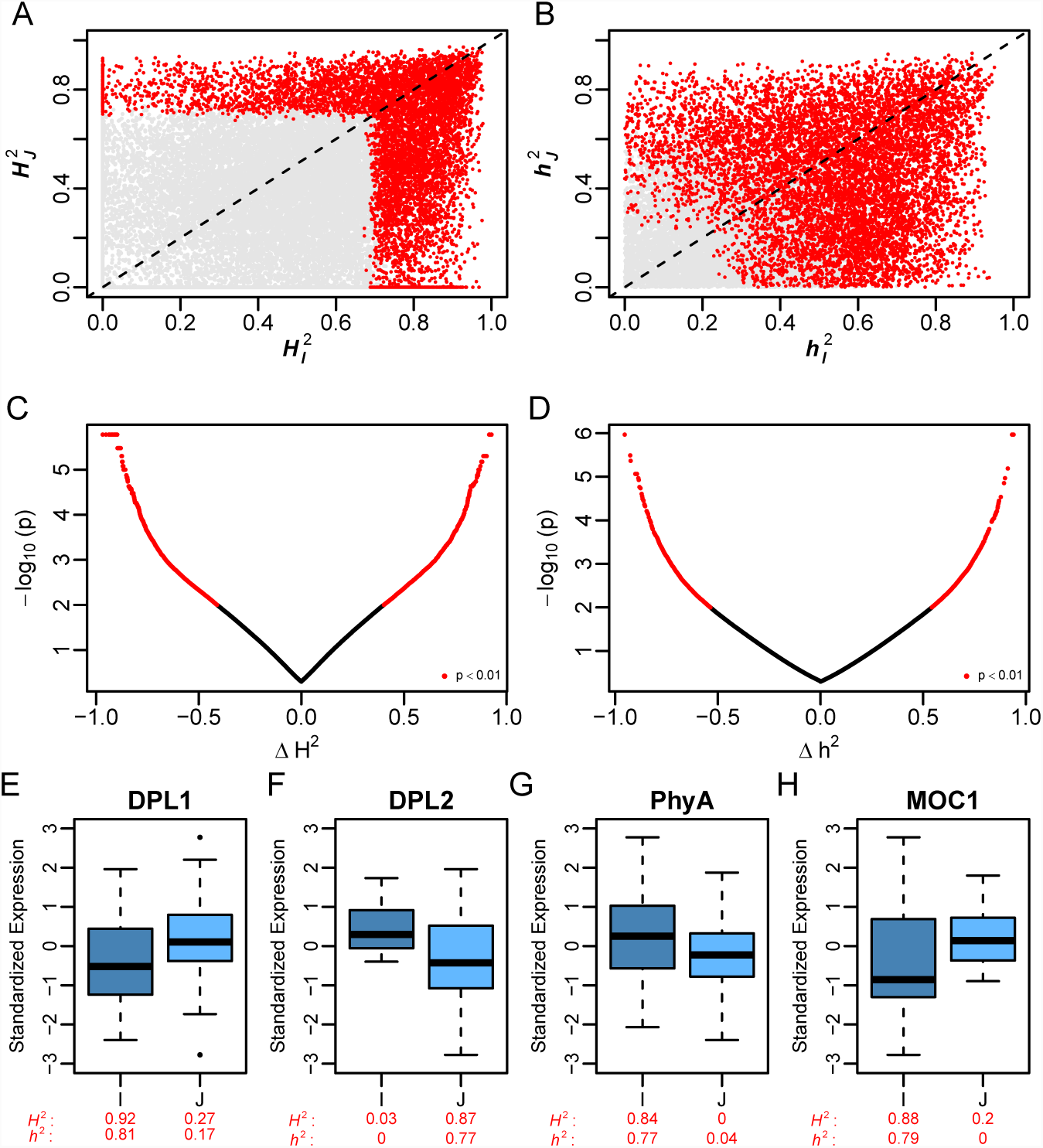
Divergent genetic variability between subspecies. (A) Comparison of broad-sense heritability between *Indica 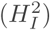* and *Japonica 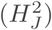*. (B) Comparisons of narrow sense heritability between the two sub-species. Red colored points in B and C indicate genes with significantly heritable expression (*FDR* < 0.001). Differences in broad (C) and narrow sense heritability (D) between *Indica* and *Japonica*. The difference in heri-tability is calculated as 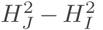 or 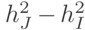. (E-H) Standardized expression of agronomically important genes showing differences in genetic variability between subspecies. The heritability is provided below each box plot. I: *Indica*, J: *Japonica*

To systematically identify genes showing significant differences in *H*^2^ or *h*^2^ (Δ*H*^2^ and Δ*h*^2^, respectively) between subspecies, accessions were randomly partitioned into two groups of equal size and the difference in heritability was estimated between groups. The resampling approach was repeated 100 times. A total of 1,860 genes showed significant differences in *H*^2^ (*p* < 0.01) between the two subspecies, with a minimum absolute difference in *H*^2^ of 0.40. Fewer genes were identified with a significant difference in *h*^2^ between *Japonica* and *Indica* (Supplemental File S4). Only 1,325 genes were found with significant differences in *h*^2^ between *Indica* and *Japonica*, and the absolute difference in *h*^2^ ranged from 0.54 to 0.95 (Figure 4).

These differences in heritability may be due to insufficient phenotypic variation (e.g. lack of expression diversity), or changes in the genetic or environmental factors that contribute to phenotypic variation. Thus, to further examine the potential causes of the observed differences in hertiability, we quantified the expression diversity (CV), genetic variation and environmental variation within each subspecies for genes exhibiting Δ*H*^2^ and Δ*h*^2^, as well as those with shared heritable variation. For genes exhibiting subspecies-specific genetic variability, the loss of heritability was largely due to an increase in environmental effects on phenotypic variation in the subspecies lacking heritability rather than loss of phenotypic variation. This is clearly evident in Supplemental Figure S2. The mean CV for Δ*H*^2^ genes decreased slightly in subspecies lacking genetic variability. However, for these same genes the proportion of phenotypic variation that was explained by environmental effects increased significantly in subspecies lacking genetic variability. Collectively, these results suggest that the differences in heritability exhibited between the subspecies is driven largely by loss of genetic variability and an increase in environmental effects rather than a loss of phenotypic variation.

Interestingly, several genes that have been reported to have divergent genetic variants between *Indica* and *Japonica* were found within Δ*H*^2^ and Δ*h*^2^ genes. For instance, *DOPPELGANGER*1 (*DPL*1) showed significantly higher *H*^2^ and *h*^2^ in *Indica* relative to *Japonica* (*H*^2^: 0.92 and 0.27, respectfully, 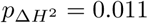; *h*^2^: 0.81 and 0.17, 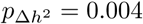; 4E). However for *DOPPELGANGER*2, the converse was true. Significantly higher *H*^2^ and *h*^2^ was observed in *Japonica* relative to *Indica* (*H*^2^: 0.87 and 0.03, 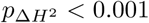; *h*^2^: 0.77 and 0, respectfully, 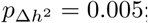; Figure 4F). Mizuta et al. (2010) showed that *DPL*1 and *DPL*2 are important regulators of *Indica-Japonica* hybrid incompatibility, and non-functional alleles arose independently for *DPL*1 and *DPL*2 within the *Indica* and *Japonica* subspecies respectively. Thus the results reported by Mizuta et al. (2010) are consistent with the divergent genetic variability in expression observed in our study. In addition to *DPL*1 and *DPL*2, a gene that is important for the regulation of shoot growth/ architecture, *MOC*1, also displayed divergent genetic variability between subspecies. *MOC*1 showed significant differences in both *H*^2^ and *h*^2^ (Figure 4H). Collectively, these results show that the two subspecies are divergent at the transcriptional and genetic levels. Moreover, many genes exhibit large differences in genetic variability between the *Indica* and *Japonica*, suggesting that these genes may be regulated by divergent genetic mechanisms.

### Joint eQTL analysis assesses cis-regulatory divergence between subspecies

The differences in the narrow-sense heritability between subspecies observed for some genes suggest a divergence in the genetic regulation of these genes. Using the transcriptional and genotypic data for this population, we next sought to identify genetic variants that could explain this divergent genetic regulation. To this end, a joint eQTL analysis was conducted across subspecies using the eQTL Bayesian model averaging (BMA) approach described by Flutre et al. (2013). With this approach, the posterior probability of specific configurations can be formally tested; in other words, the probability that an eQTL is present/active in both the *Indica* and *Japonica* subspecies or unique to a given subspecies can be determined. The 91 accessions were classified into *Indica* and *Japonica* subspecies using STRUCTURE as described earlier, yielding 35 *Indica*-type and 56 *Japonica*-type accessions. eQTLs were modeled for genes showing significant *H*^2^ in at least one subspecies (6,307 genes) and 274,499 SNPs. For each gene, associations were tested for SNPs within 100kb of the transcription start site. A total of 5,097 genes were detected with one or more eQTL at an FDR of 0.05 (Supplemental File S5). This equates to approximately 81% of the genes displaying heritable expression, and indicates that a large portion of genes with heritable expression are regulated by variants in close proximity to the gene.

To identify eQTL genes that were specific to a given subspecies, the SNP with the highest probability of being the eQTL was selected for each gene, and the posterior probability for all three configurations (*Indica*-specific, *Japonica*-specific, and across subspecies) was compared. Of the 5,097 eQTL genes detected, 80% (4,077 genes; 3,826 unique SNPs) were detected across subspecies, 18% (914 genes; 880 unique SNPs) were detected for *Indica* accessions, and 2% (106 genes; 103 unique SNPs) were detected only in *Japonica* accessions. These results indicate that while a large portion of *cis*-eQTLs are shared across the two subspecies of cultivated rice, many genes are regulated by unique *cis* regulatory mechanisms that are specific to the *Indica* subspecies.

### Signatures of selection are evident among subspecies specific eQTL

The presence or absence of cis-regulatory variants within a given subspecies may be the result of the unique domestication histories that have shaped *Indica* and *Japonica*, and/or driven by environmental adaptation of the wild progenitors from which they were derived. The absence of variation at the eQTL SNP could be due to sampling during differentiation of the wild progenitors or during domestication (e.g. lost purely by chance), or due to selective pressures imposed by the environment or humans. In the case of selection, we expect to see reduced genetic diversity around the eQTL compared to the rest of the genome. To determine whether the absence of subspecies-specific eQTL are the result of selection, we calculated the average nucleotide diversity (*π*) in 100 Kb windows around significant subspecies-specific eQTL within each subspecies and compared these values to the overall average *π* for 100 Kb windows across the genome within each subspecies using a two-sided *t*-test. Comparisons within each subspecies of *π* for eQTLs and the genome-wide average should account for the inherent differences in *π* between the two subspecies.

**Figure 5.**
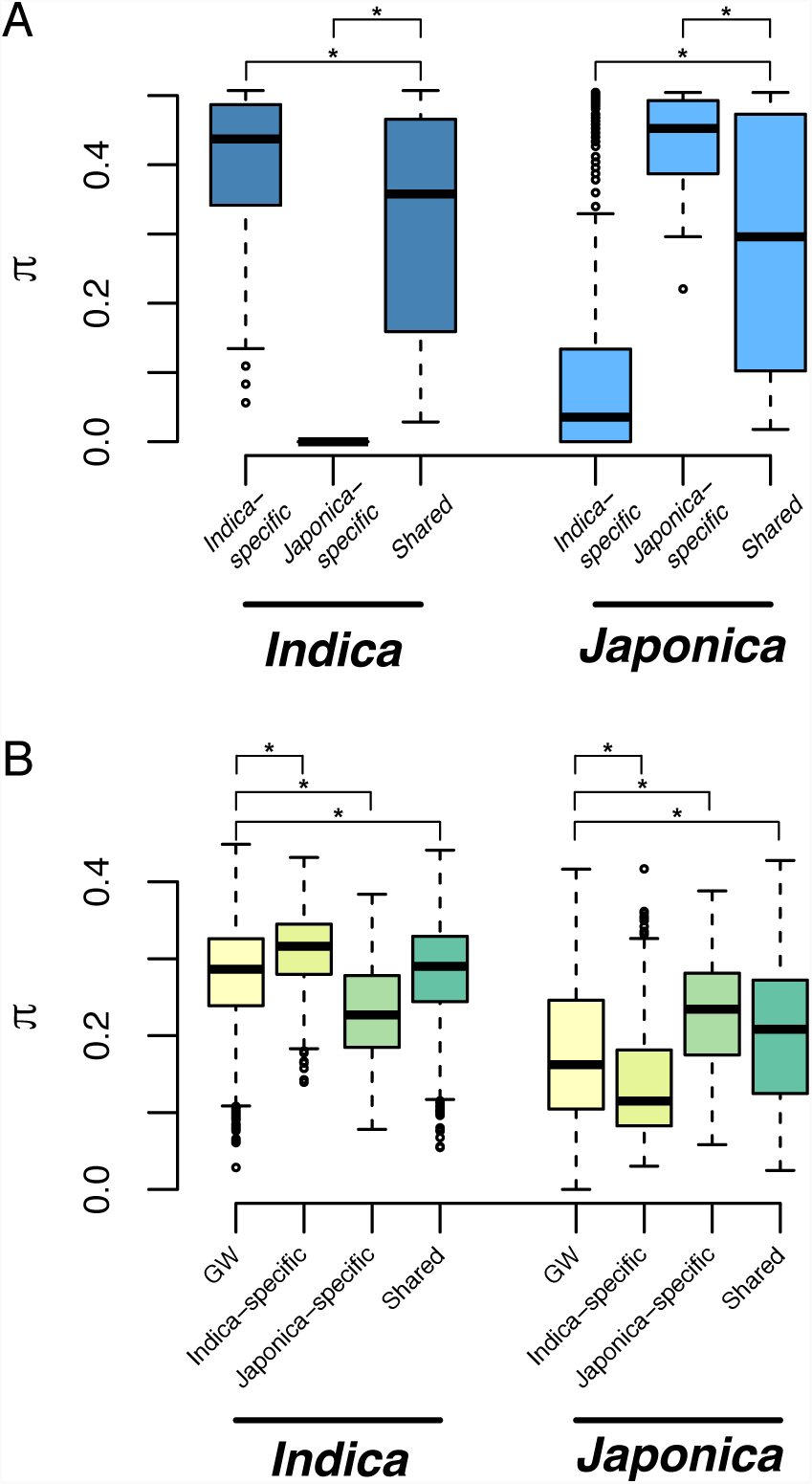
Nucleotide diversity at cis-eQTL. (A) Nucleotide diversity (*π*) for the most significant SNP for each cis-eQTL. The distribution of *π* is pictured fro each subspecies and each eQTL type. (B) Distribution of *π* for 100 Kb windows around the most significant SNP for each cis-eQTL. Genome-wide (GW) *π* was determined by randomly selecting × SNPs that were more than 100 kb from a cis-eQTL and low diversity SNPs (MAF *<* 0.1 in both subspecies) were removed prior to analyses. Asterisks indicate a significant differences determined via Tukey’s test between eQTL types (*p* < 1 × 10^-8^).

Consistent with what would be expected under selection, a significant reduction in nucleotide diversity was observed for eQTL SNPs that were absent in a subspecies, as well as for regions around subspecies-specific eQTL (Figure 3). For instance, for *Indica*-specific eQTL, the average *π* in *Japonica* was approximately 22% lower than the genome-wide average (0.138 and 0.176, respectively; *p* < 1 × 10^-15^). Similarly, the average *π* in *Indica* for *Japonica*-specific eQTL was about 16% lower than the genome-wide average (0.235 and 0.279, respectively; *p* = 3.85 × 10^-10^). Interestingly, slightly higher nucleotide diversity was observed for regions around subspecies-specific eQTL in subspecies in which they were detected compared to genome-wide nucleotide diversity, as well as for shared eQTL when compared to genome-wide nucleotide diversity. Collectively, these results indicate that the absence of eQTL within a given subspecies may be the result of selective pressures that reduced genetic diversity within the eQTL regions.

Given the small sample size in the current study (*n* = 91) we sought to confirm these results using resequencing data for a larger population of 3,024 diverse rice accessions (Wang et al., 2018; Mansueto et al., 2016a,b; Alexandrov et al., 2014). To this end, we extracted SNP information for 3,024 rice accessions in the same 100 Kb window surrounding eQTL, and examined *π* within each subpopulation for these regions. As above, *π* within these regions were compared with genome-wide averages for 100 kb windows. The 3,024 rice accessions are classified into 12 subpopulations: *admix* (103 accessions), *aromatic* (76 accessions), *aus* (201 accessions), *indica1A* (209 accessions), *indica1B* (205 accessions), *indica2* (285 accessions), *indica3* (475 accessions), *indica-X* (615 accessions), *japonica-X* (83 accessions), *subtropical japonica* (112 accessions), *temperate japonica* (288 accessions), and *tropical japonica* (372 accessions). The *Indica* subspecies are represented by *indica1A*, *indica1B*, *indica2*, *indica3*, and *indica-X*; while the *Japonica* subspecies consists of the *japonica-X*, *subtropical japonica*, *temperate japonica*, and *tropical japonica* subpopulations.

Consistent with the results derived from the 91 accessions, *π* within subspecies-specific eQTL was lower in subpopulations lacking the eQTL. For instance, for the *Japonica* subpopulations (*japonica-x*, *subtropical japonica*, *temperate japonica*, and *tropical japonica*) *π* estimates for *Indica*-specific eQTL were considerably lower than those for *Indica* subpopulations (*indica-1A*, *indica-1B*, *indica-2*, *indica-3*, and *indica-x*). The converse was true for *Japonica*-specific eQTL, with lower *π* observed in *Indica* subpopulations relative to *Japonica*. However for the shared eQTL, *π* estimates were higher than the genome-wide averages, suggesting that genetic diversity within regions that regulate gene expression is maintained.

To identify specific loci that may have been targeted by selection, we selected eQTL regions with an average *π* within a 100 Kb window that was below the the 5% quantile for genome-wide average for a given subspecies. Consistent with the results above, we observed a greater frequency of low diversity eQTL regions in subspecies lacking the subspecies-specific eQTL. For instance, approximately 11% of the 880 *Indica*-specific eQTL were found in regions of low diversity in *Japonica* (*π*_*Jap*_ *≤* 0.0645). While for *Japonica*-specific eQTL, 14% (14 of the 103) eQTL regions were lying in regions of low diversity in *Indica* (*π*_*Ind*_ *≤* 0.1617). However, for shared eQTL and for subspecies in which the subspecies-specific eQTL was detected, the converse was true. Only a small percentage of eQTL regions were found within regions of low diversity. For instance, approximately 3.5% of shared eQTL were found in regions of low diversity in both *Indica* and *Japonica*, and less than 1% of subspecies eQTL were found in regions of low diversity in the subspecies in which they were detected. Collectively these results suggest that selective pressures may have shaped the cis-regulatory divergence of the *Indica* and *Japonica* subspecies.

**Figure 6.**
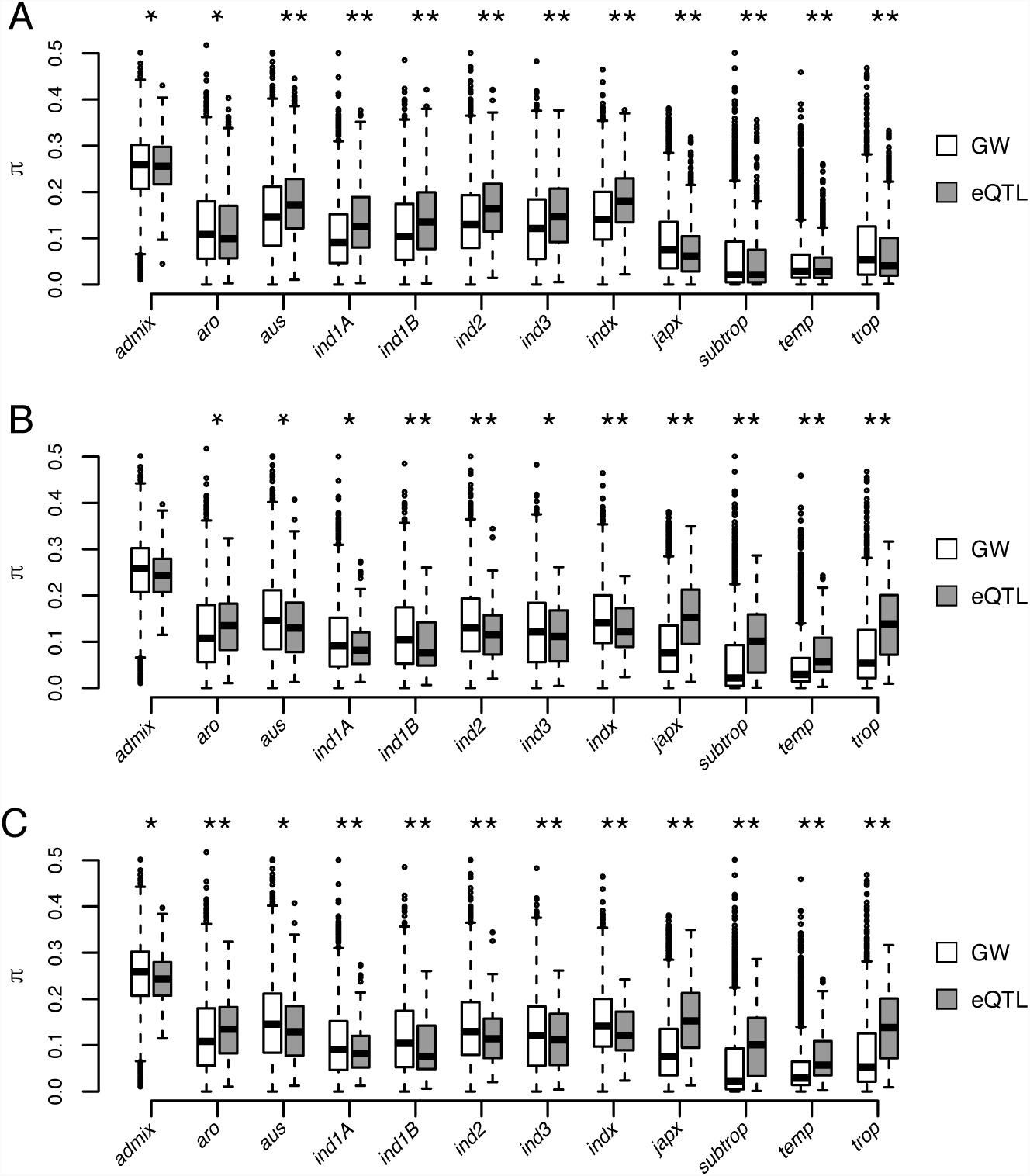
Nucleotide diversity at cis-eQTL within subpopulations for 3,053 rice accessions. Average nucleotide diversity (*π*) for 100 kb regions surrounding *Indica*-specific, *Japonica*-specific, and shared eQTL are pictured in panels A, B, and C, respectively. For each, subpopulation and class of eQTL (e.g.*Indica*-specific, *Japonica*-specific, and shared) *π* was calculated for each SNP within 100 kb of the most significant eQTL SNP. *π* for the eQTL windows were compared to a genome wide (GW) average in which regions with eQTL and site with low diversity (MAF *<* 0.01 in 10 of 12 subpopulations) were excluded. Asterisks indicate significant differences between GW and eQTL regions determined using a two-sided Student’s *t*-test (* *p* < 0.05; ** *p* < 0.001). Subpopulations are named following Wang et al. (2018) (*aro*: aromatic; *ind1A*: indica-1A; *ind1B*: indica-1B; *ind2*: indica-2; *indx*: indica-X; *japx*: japonica-X; *subtrop*: subtropical japonica; *temp*: temperate japonica; *trop*: tropical japonica).

## Discussion

The differentiation between the *Indica* and *Japonica* subspecies of cultivated rice has been intensively studied at the morphological, biochemical, and genetic levels (Kato, 1928; Terao and Mizushima, 1942; Matsuo, 1952; Morinaga, 1954; Morishima and Oka, 1981; Glaszmann, 1987; Goff et al., 2002; Yu et al., 2002; Feltus et al., 2004; Stein et al., 2018; Koide et al., 2018; Schatz et al., 2014; Huang et al., 2008; Wang et al., 2014; Huang et al., 2012b). However, the divergence at the transcriptional levels remains understudied. Here, we provide a comprehensive analysis of the transcriptional and *cis*-regulatory divergence between the major subspecies of rice, and show that the presence or absence of cis regulatory variants within the subspecies is a component of this divergence.

The transcriptional divergence is most evident in the large number of expressed genes showing differences in the magnitude or frequency of expression. Of the 25,732 genes showing evidence of expression in the current study, approximately 29% showed significant differences in expression levels between the two subspecies. Moreover, approximately 8% of expressed genes showed evidence of presence-absence expression variation. While few studies have examined the differences in expression levels between diverse populations of *Indica* and *Japonica*, recent studies have utilized whole genome sequencing to shed light on the genetic differentiation between the subspecies of cultivated rice (Huang et al., 2012b; Wang et al., 2018). In a recent study, Wang et al. (2018) found that on average approximately 15% of all genes showed evidence of PAV between the genomes of *Indica* and *Japonica* accessions, further indicating that PAV is pervasive between the subspecies of cultivated rice. While the number of PAV reported by Wang et al. (2018) are nearly two fold higher than those reported in the current study, it is important to note that only a single tissue was sampled for 91 accessions at a single time point. Therefore, while the expression data provides considerable insight into transcriptional variation in cultivated rice, it likely captures only a portion of the total transcriptome given the lack of temporal and spatial resolution. Moreover, Wang et al. (2018) captured PAV using 3,010 resequenced rice genomes, while the current study utilized only a fraction of the variation of Wang et al. (2018) with RNA sequencing of 91 accessions. Thus, increased sample size via larger populations and more sampling within tissue and developmental context may lead to a better agreement between PAV at the genome and transcriptional levels.

### Potential causes of transcriptional divergence between *Indica* and *Japonica*

Lower mean expression values or absence of expression in a given subspecies may be the result of both heritable and non-heritable effects. The availability of high density SNP information for RDP1 allowed us to begin to elucidate the genetic basis of the observed transcriptional divergence between the subspecies of cultivated rice. A notable portion of genes with evidence of PAV or DE also showed differences in genetic variability between the subspecies (13% and 9% of DE genes showed differences in *H*^2^ and *h*^2^, respectively, and 20% and 15% of PAV genes showed differences in *H*^2^ and *h*^2^), indicating that for many genes, the genetic mechanisms that regulate expression may be different between the two subspecies. However, many genes that display divergent expression patterns have non-significant differences in genetic variability. There are several explanations for this. For one, the thresholds used to identify genetically divergent genes were quite stringent. For instance, genes must have a difference in genetic variability in either the broad sense greater than 0.4022 between subspecies to be labeled as statistically significant, and in the narrow sense 0.5364. Therefore, it is possible that many more DE or PAV genes have different genetic architectures in the two subspecies, but were missed because of the statistical threshold selected. A second possibility is that many of the genes showed divergent expression are influenced greatly by the environment, and thus have low heritability. Thus, these genes would be filtered out in these genetic analyses.

The heritable transcriptional divergence may be due to genetic variants that influence gene expression and are divergent between *Indica* and *Japonica*. These include large structural variants (e.g. deletions, insertions, inversions, and/or duplications), or SNPs that may act in cis or trans to influence gene regulation. While high density SNP information is available for this population and can be leveraged to identify SNPs that regulate expression and are divergent between the subspecies, the identification of larger structural variants that influence expression is only attainable through full genome sequencing, which is not currently available for RDP1. As more genetic resources become available for RDP1 this would be a promising future direction to resolve the causal basis of these transcriptional differences.

The availability of high density SNP information for RDP1 allowed us to begin to elucidate the genetic basis of the observed transcriptional divergence between the subspecies of cultivated rice, and classify genetic effects into those that are common between subspecies, or unique to a given subspecies. While the eQTL-BMA approach has proven to be a powerful framework for assessing the specificity of eQTL for a given tissue or population, one potential limitation of eQTL-BMA is that the framework only allows modeling cis-eQTL. Trans-eQTLs are often difficult to detect due the penalties associated with the large number of statistical tests performed, and because trans-eQTL often have small effect sizes and thus require larger dataset for detection. Several studies in humans have shown that cis-eQTL typically only explain 30-40% of genetic variation in expression (Price et al., 2011; Grundberg et al., 2012; Hore et al., 2016). Thus, the divergent regulatory variants captured in the current study only reflect a portion of the differences in genetic variation between the two subspecies. Further studies are necessary to shed light on the contribution of trans-regulatory variants on the genetic differentiation between *Indica* and *Japonica* transcriptomes.

The joint eQTL analysis facilitated the identification of 5,097 genes associated with one or more SNP in *cis*. For most of these genes (81%), the cis-regulatory variant was shared between both subspecies, indicating that much of the cis-regulatory variation is common between the two subspecies. This high degree of overlap is somewhat expected. For one, both *Indica* and *Japonica* originate from populations of the same species, *Oryza rufipogon*. Moreover, crosses between *Indica* and *Japonica* often produce viable offspring, indicating a high degree of colinearity and functional similarity between the genomes. Thus, while considerable differentiation between founder *Oryza rufipogon* populations has been reported and further divergence has likely occurred since domestication, the common origin and inter-specific comparability suggests that the transcriptional regulation and genome structure is similar (Huang et al., 2012b).

Despite the majority of cis-regulatory variants being shared between the two subspecies, approximately 18% of all genes with one or more eQTL were found to be unique to *Indica* or *Japonica*. The large majority of these subspecies-specific eQTL were detected in the *Indica* subspecies and were nearly fixed in *Japonica* indicating low genetic diversity at the eQTL. Moreover, the genetic variation surrounding subspecies-specific eQTL were significantly lower that genome wide averages, indicating that selective pressures may have uniquely shaped components of *cis*-regulatory variation between the two subspecies. The two subspecies are derived from geographically and genetically distinct subpopulations of *Oryza rufipogon* (Huang et al., 2012b). Therefore, it remains an open question whether these events occurred during the differentiation between *O. rufipogon* subpopulations or during the domestication of *O. sativa*.

We found significantly higher nucleotide diversity in the regions surrounding eQTL compared to genome wide averages. These patterns of diversity were consistent within subpopulations for shared eQTL, as well as for subspecies-specific eQTL in the subspecies or subpopulations in which they were detected. Although the functions for the majority of these eQTL genes are unknown, the observation that their expression is regulated at a genetic level suggests that they may play a role in the regulation of some biological process. Genetic diversity is a prerequisite to evolutionary change (Lewontin et al., 1974). Therefore the higher nucleotide diversity at these regions compared to genome-wide backgrounds may be reflective of the importance of maintaining genetic variation for these biological processes through regulation at the transcriptional level.

### Functional significance of transcriptional divergence

The current study sheds light on the transcription divergence between the major subspecies of cultivated rice. Many of these genes found to have divergent expression, genetic variability, or regulatory variation have been reported to be underlying important agronomic traits, such as photoperiod adaptation and development. Therefore these observed differences may have potential agronomic significance.

Among these divergent genes, we identified three genes (*OsPhyA, OsPhyC, and OsCO3*), that have been reported to be associated with the timing of reproductive development in response to day length that had significant heritability in *Indica* only. The two phytochrome genes, *OsPhyA and OsPhyC* are activated under long-day conditions and repress flowering time through *OsGhd7* (Takano et al., 2005; Lee et al., 2016).

Although no studies have shown whether *OsCO3* participate directly in the pathway involving *OsPhy* genes, disruption of *OsCO3* interferes with photoperiod sensitivity and/or flowering time (Kim et al., 2008). For instance, Kim et al. (2008) showed that the overexpression of *OsCO3* delayed flowering under short-day conditions. In most rice varieties, short-days promote the transition from vegetative to reproductive growth (Song et al., 2015). However, *temperate japonica* rice varieties adapted to higher latitudes have been selected to initiate flowering in long-days to escape the negative impact of low temperatures in autumn on pollen fertility (Huang et al., 2012a; Itoh et al., 2004; Naranjo et al., 2014). All genes showed heritable expression only in the *Indica* subspecies, indicating that in the *Japonica* subspecies expression variation may be driven largely by non-genetic effects. Moreover, the patterns of genetic variability for these genes are consistent with their potential role in the adaptation of flowering in different environments for *Indica* and *Japonica*.

In addition to genes regulating phenology, several genes were identified that have been reported to play important roles in the regulation of shoot architecture (*D*18, *MT* 2*b*, and *MOC*1). For instance, two genes *dwarf* 18 (*D*18) and *Metallothionein*2*b* (*MT* 2*b*) have been reported to regulate plant height (Itoh et al., 2001; Yuan et al., 2008). *D*18 encodes a GA-*β* hydroxylase and is involved with GA biosynthesis. Loss of function mutants exhibit a severe dwarf phenotype (Itoh et al., 2001). Interestingly, *D*18 was found have an *Indica*-specific eQTL, but did not exhibit a difference in *H*^2^ or *h*^2^ between the two subspecies (*p* = 0.046 and *p* = 0.19, respectively), indicating that genetic differences may be confined to local regions around *D*18. The diversity within the 100kb regions surrounding the eQTL region was quite high compared to the genome-wide average in both subspecies (*π*_*Ind*_ = 0.27, *π*_*Jap*_ = 0.18) indicating that the absence of the *D*18 eQTL in *Japonica* may be due to low diversity within the eQTL SNP, rather than potential selective pressures between subspecies.

## Conclusions

The morphological and genetic differences between subspecies of cultivated rice have been studied extensively, however the divergence of *Indica* and *Japonica* at the transcriptional and regulatory levels is largely unresolved. Here, we provide, to date, the first detailed population-level characterization of transcriptional diversity within cultivated rice, and assess the divergence in trancriptomes and expression variation between *Indica* and *Japonica*. We find that many agronomically important genes exhibit differences in expression levels, and/or cis-regulatory variation between the subspecies. These resources provided by this study can serve as a foundation for future functional genomics studies in rice, and can be further utilized to connect gene function with natural variation in gene expression.

## Acknowledgments

Funding for this research was provided by the National Science Foundation (United States) through Awards 1238125 and 1736192 to Harkamal Walia.

## Data Availability

All transcriptional data can be accessed via NCBI Gene Expression Omnibus under accession number GSE98455.

## Supplemental Data

**Table S1.**
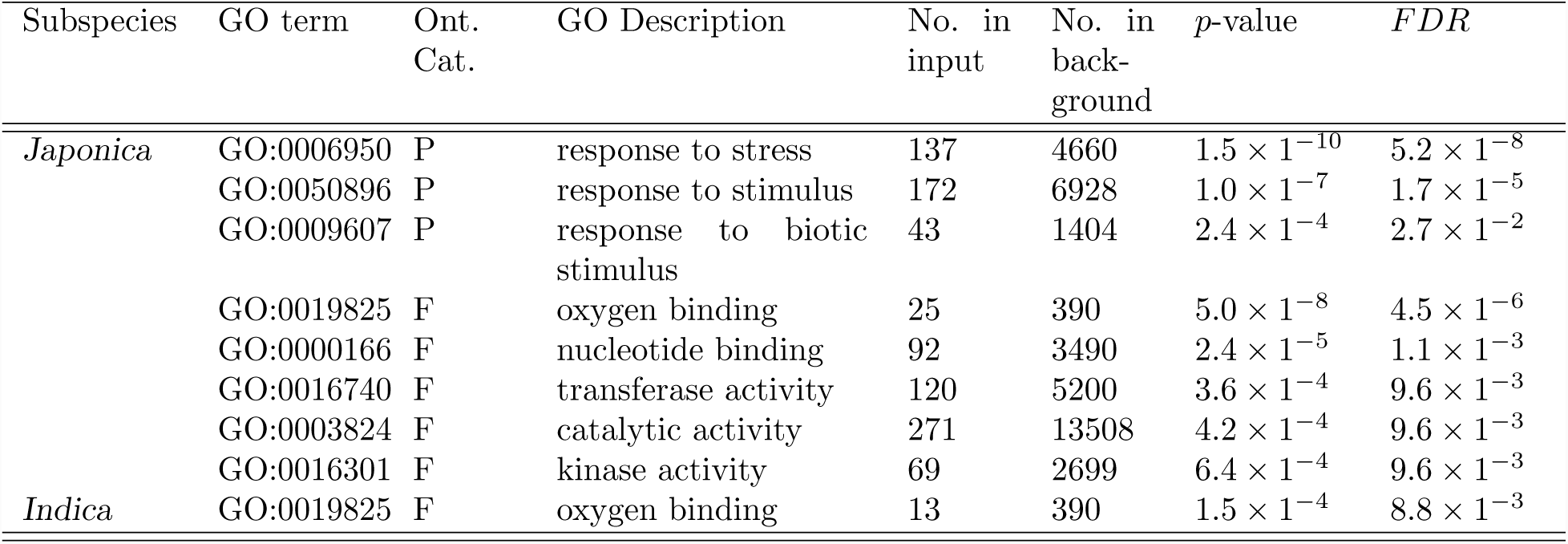
Gene onotology (GO) enrichment analysis for genes exhibiting significant presence-absence expression variation (PAV) (*FDR* < 0.001). GO enrichment was conducted using AgriGO (http://bioinfo.cau.edu.cn/agriGO) using the MSU V7 genome build without transposable elements as a background. GO enrichment was conducted separately for genes expressed predominately in each subspecies.

| Subspecies | GO term | Ont.<br>Cat. | GO Description | No. in<br>input | No. in<br>back-<br>ground | $p$-value | $FDR$ |
| --- | --- | --- | --- | --- | --- | --- | --- |
| *Japonica* | GO:0006950 | P | response to stress | 137 | 4660 | $1.5 \times 10^{-10}$ | $5.2 \times 10^{-8}$ |
| | GO:0050896 | P | response to stimulus | 172 | 6928 | $1.0 \times 10^{-7}$ | $1.7 \times 10^{-5}$ |
| | GO:0009607 | P | response to biotic stimulus | 43 | 1404 | $2.4 \times 10^{-4}$ | $2.7 \times 10^{-2}$ |
| | GO:0019825 | F | oxygen binding | 25 | 390 | $5.0 \times 10^{-8}$ | $4.5 \times 10^{-6}$ |
| | GO:0000166 | F | nucleotide binding | 92 | 3490 | $2.4 \times 10^{-5}$ | $1.1 \times 10^{-3}$ |
| | GO:0016740 | F | transferase activity | 120 | 5200 | $3.6 \times 10^{-4}$ | $9.6 \times 10^{-3}$ |
| | GO:0003824 | F | catalytic activity | 271 | 13508 | $4.2 \times 10^{-4}$ | $9.6 \times 10^{-3}$ |
| | GO:0016301 | F | kinase activity | 69 | 2699 | $6.4 \times 10^{-4}$ | $9.6 \times 10^{-3}$ |
| *Indica* | GO:0019825 | F | oxygen binding | 13 | 390 | $1.5 \times 10^{-4}$ | $8.8 \times 10^{-3}$ |

**Figure S1.**
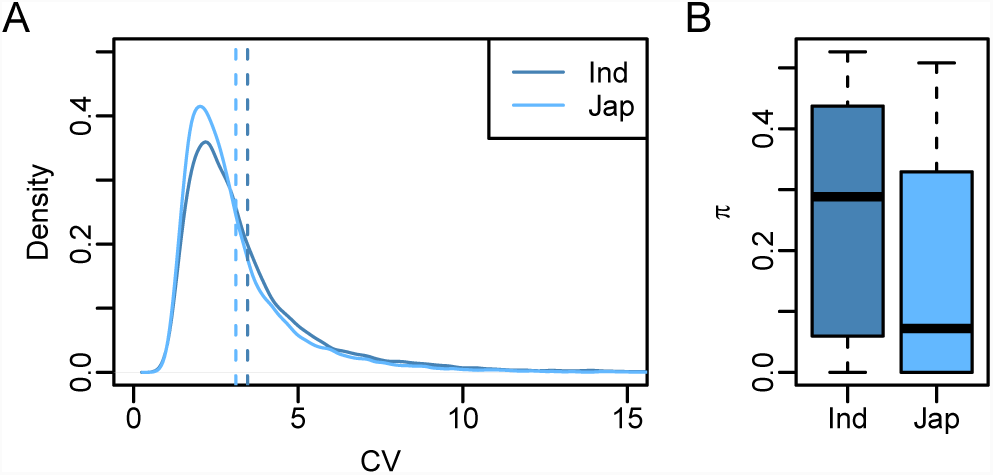
Genetic and expression diversity within *Indica* and *Japonica* accessions. (A) The coefficient of variation was used as an estimate of the diversity in gene expression within each subspecies. The vertical dashed lines represent the mean CV within each subspecies. (B) Site-wise nucleotide diversity (*π*) was used as an estimate of the genetic diversity within each of the subspecies using 36,901 SNPs described by Zhao et al. (2011).

**Figure S2.**
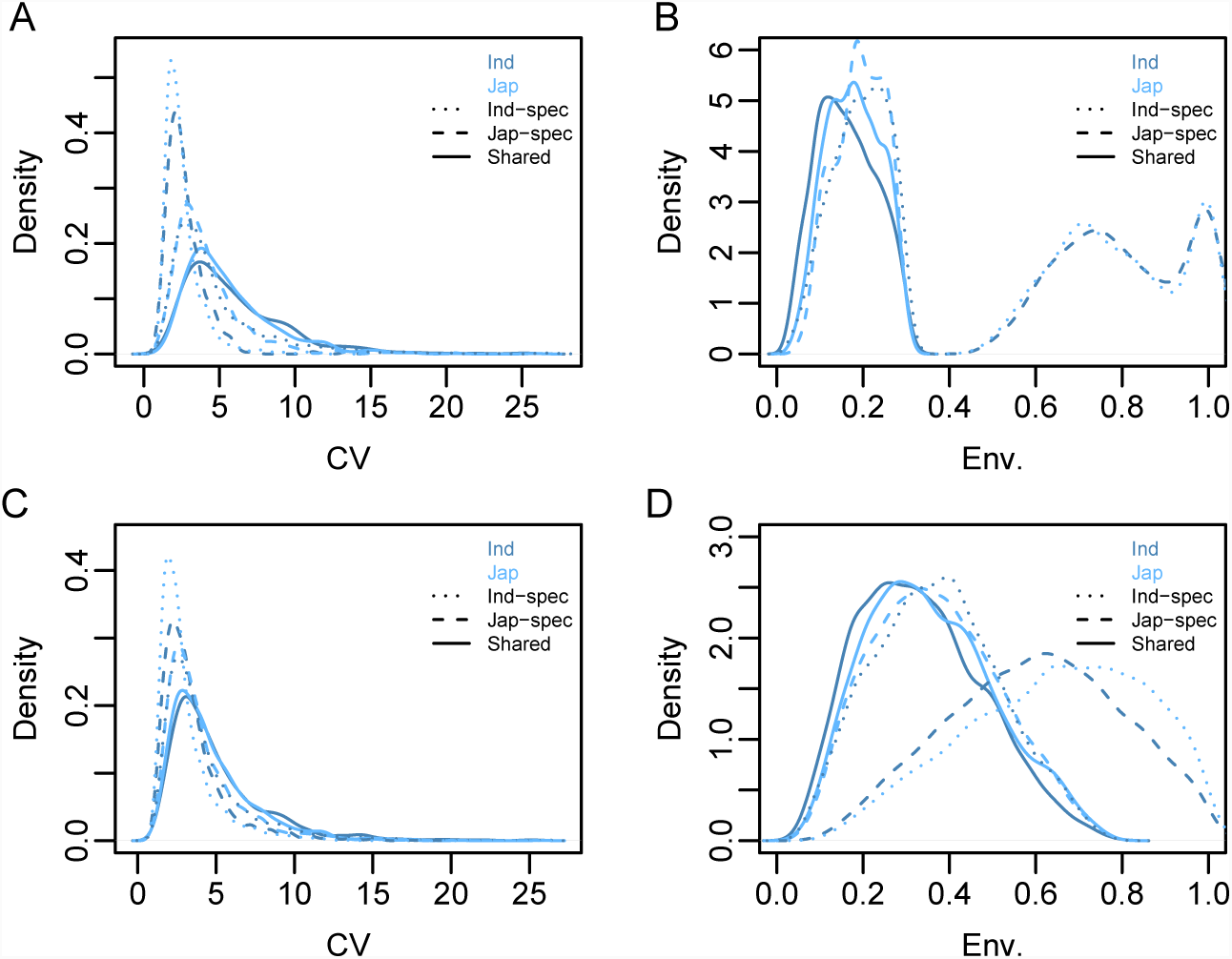
Assessing phenotypic variation and environmental effects for genes exhibiting genetic variability within each sub-species. Genes were classified into three categories based on their patterns of genetic variability.” Shared” refers to genes showing significant genetic variability (*FDR* < 0.001) in both subspecies. The categories” *Indica*- specific” and” *Japonica*-specific” refer to genes that showed significant differences in genetic variability (e.g. Δ*H*^2^ or Δ*h*^2^) and had heritable expression in *Indica* and *Japonica*, respectively. Phenotypic variation was assessed using the coefficient of variation (CV) for *H*^2^ or *h*^2^ genes (A and C, respectively). The contribution of the environment on phenotypic variation was determined as 1 − *H*^2^ and 1 − *h*^2^ (B and D, respectively). The categories of genetic variability are indicated by line type, while the subspecies in which CV or environmental variation was measured are indicated by the color of lines.

## References

Alexandrov, N., Tai, S., Wang, W., Mansueto, L., Palis, K., Fuentes, R. R., Ulat, V. J., Chebotarov, D., Zhang, G., Li, Z., et al. (2014). Snp-seek database of snps derived from 3000 rice genomes. Nucleic acids research, 43(D1):D1023–D1027.

Anders, S., Pyl, P. T., and Huber, W. (2015). Htseq—a python framework to work with high-throughput sequencing data. Bioinformatics, 31(2):166–169.

Andrews, S. et al. (2010). FastQC: a quality control tool for high throughput sequence data.

Bates, D., Mächler, M., Bolker, B., and Walker, S. (2015). Fitting linear mixed-effects models using lme4. Journal of Statistical Software, 67(1):1–48.

Benjamini, Y. and Hochberg, Y. (1995). Controlling the false discovery rate: a practical and powerful approach to multiple testing. Journal of the royal statistical society. Series B (Methodological), pages 289–300.

Bolger, A. M., Lohse, M., and Usadel, B. (2014). Trimmomatic: a flexible trimmer for illumina sequence data. Bioinformatics, 30(15):2114–2120.

Butler, D., Cullis, B. R., Gilmour, A., and Gogel, B. (2009). Asreml-r reference manual. The State of Queensland, Department of Primary Industries and Fisheries, Brisbane.

Caicedo, A. L., Williamson, S. H., Hernandez, R. D., Boyko, A., Fledel-Alon, A., York, T. L., Polato, N. R., Olsen, K. M., Nielsen, R., McCouch, S. R., et al. (2007). Genome-wide patterns of nucleotide polymorphism in domesticated rice. PLoS genetics, 3(9):e163.

Danecek, P., Auton, A., Abecasis, G., Albers, C. A., Banks, E., DePristo, M. A., Handsaker, R. E., Lunter, G., Marth, G. T., Sherry, S. T., et al. (2011). The variant call format and vcftools. Bioinformatics, 27(15):2156–2158.

Ding, J., Araki, H., Wang, Q., Zhang, P., Yang, S., Chen, J.-Q., and Tian, D. (2007). Highly asymmetric rice genomes. BMC genomics, 8(1):154.

Eizenga, G. C., Ali, M., Bryant, R. J., Yeater, K. M., McClung, A. M., McCouch, S. R., et al. (2014). Registration of the rice diversity panel 1 for genomewide association studies. Journal of Plant Registrations, 8(1):109–116.

Famoso, A. N., Zhao, K., Clark, R. T., Tung, C.-W., Wright, M. H., Bustamante, C., Kochian, L. V., and McCouch, S. R. (2011). Genetic architecture of aluminum tolerance in rice (Oryza sativa) determined through genome-wide association analysis and QTL mapping. PLoS genetics, 7(8):e1002221.

Feltus, F. A., Wan, J., Schulze, S. R., Estill, J. C., Jiang, N., and Paterson, A. H. (2004). An snp resource for rice genetics and breeding based on subspecies indica and japonica genome alignments. Genome research, 14(9):1812–1819.

Flutre, T., Wen, X., Pritchard, J., and Stephens, M. (2013). A statistical framework for joint eqtl analysis in multiple tissues. PLoS genetics, 9(5):e1003486.

Garris, A. J., Tai, T. H., Coburn, J., Kresovich, S., and McCouch, S. (2005). Genetic structure and diversity in Oryza sativa L. Genetics, 169(3):1631–1638.

Glaszmann, J.-C. (1987). Isozymes and classification of asian rice varieties. Theoretical and Applied Genetics, 74(1):21–30.

Goff, S. A., Ricke, D., Lan, T.-H., Presting, G., Wang, R., Dunn, M., Glazebrook, J., Sessions, A., Oeller, P., Varma, H., et al. (2002). A draft sequence of the rice genome (Oryza sativa L. ssp. japonica). Science, 296(5565):92–100.

Grundberg, E., Small, K. S., Hedman, A. K., Nica, A. C., Buil, A., Keildson, S., Bell, J. T., Yang, T.-P., Meduri, E., Barrett, A., et al. (2012). Mapping cis-and trans-regulatory effects across multiple tissues in twins. Nature genetics, 44(10):1084.

Hore, V., Vinũela, A., Buil, A., Knight, J., McCarthy, M. I., Small, K., and Marchini, J. (2016). Tensor decomposition for multiple-tissue gene expression experiments. Nature genetics, 48(9):1094.

Huang, C.-L., Hung, C.-Y., Chiang, Y.-C., Hwang, C.-C., Hsu, T.-W., Huang, C.-C., Hung, K.-H., Tsai, K.-C., Wang, K.-H., Osada, N., et al. (2012a). Footprints of natural and artificial selection for photoperiod pathway genes in Oryza. The Plant Journal, 70(5):769–782.

Huang, X., Kurata, N., Wang, Z.-X., Wang, A., Zhao, Q., Zhao, Y., Liu, K., Lu, H., Li, W., Guo, Y., et al. (2012b). A map of rice genome variation reveals the origin of cultivated rice. Nature, 490(7421):497.

Huang, X., Lu, G., Zhao, Q., Liu, X., and Han, B. (2008). Genome-wide analysis of transposon insertion polymorphisms reveals intraspecific variation in cultivated rice. Plant physiology, 148(1):25–40.

Huang, X., Sang, T., Zhao, Q., Feng, Q., Zhao, Y., Li, C., Zhu, C., Lu, T., Zhang, Z., Li, M., et al. (2010). Genome-wide association studies of 14 agronomic traits in rice landraces. Nature genetics, 42(11):961.

Itoh, H., Tatsumi, T., Sakamoto, T., Otomo, K., Toyomasu, T., Kitano, H., Ashikari, M., Ichihara, S., and Matsuoka, M. (2004). A rice semi-dwarf gene, tan-ginbozu (d35), encodes the gibberellin biosynthesis enzyme, ent-kaurene oxidase. Plant molecular biology, 54(4):533–547.

Itoh, H., Ueguchi-Tanaka, M., Sentoku, N., Kitano, H., Matsuoka, M., and Kobayashi, M. (2001). Cloning and functional analysis of two gibberellin 3β-hydroxylase genes that are differently expressed during the growth of rice. Proceedings of the National Academy of Sciences, 98(15):8909–8914.

Jung, K.-H., Gho, H.-J., Giong, H.-K., Chandran, A. K. N., Nguyen, Q.-N., Choi, H., Zhang, T., Wang, W., Kim, J.-H., Choi, H.-K., et al. (2013). Genome-wide identification and analysis of japonica and indica cultivar-preferred transcripts in rice using 983 affymetrix array data. Rice, 6(1):19.

Kato, A. (1928). On the affinity of rice varieties as shown by the fertility of rice plants. Centr. Agric. Inst. Kyushu Imp. Univ., 2:241–276.

Kim, S.-K., Yun, C.-H., Lee, J. H., Jang, Y. H., Park, H.-Y., and Kim, J.-K. (2008). Osco3, a constans-like gene, controls flowering by negatively regulating the expression of ft-like genes under sd conditions in rice. Planta, 228(2):355.

Koide, Y., Ogino, A., Yoshikawa, T., Kitashima, Y., Saito, N., Kanaoka, Y., Onishi, K., Yoshitake, Y., Tsukiyama, T., Saito, H., et al. (2018). Lineage-specific gene acquisition or loss is involved in interspecific hybrid sterility in rice. Proceedings of the National Academy of Sciences, 115(9):E1955–E1962.

Lee, Y.-S., Yi, J., and An, G. (2016). Osphya modulates rice flowering time mainly through osgi under short days and ghd7 under long days in the absence of phytochrome b. Plant molecular biology, 91(4-5):413–427.

Lewontin, R. C. et al. (1974). The genetic basis of evolutionary change, volume 560. Columbia University Press New York.

Love, M. I., Huber, W., and Anders, S. (2014). Moderated estimation of fold change and dispersion for rna-seq data with deseq2. Genome biology, 15(12):550.

Lu, T., Lu, G., Fan, D., Zhu, C., Li, W., Zhao, Q., Feng, Q., Zhao, Y., Guo, Y., Li, W., et al. (2010). Function annotation of rice transcriptome at single nucleotide resolution by rna-seq. Genome research, pages gr–106120.

Mansueto, L., Fuentes, R. R., Borja, F. N., Detras, J., Abriol-Santos, J. M., Chebotarov, D., Sanciangco, M., Palis, K., Copetti, D., Poliakov, A., et al. (2016a). Rice snp-seek database update: new snps, indels, and queries. Nucleic acids research, 45(D1):D1075–D1081.

Mansueto, L., Fuentes, R. R., Chebotarov, D., Borja, F. N., Detras, J., Abriol-Santos, J. M., Palis, K., Poliakov, A., Dubchak, I., Solovyev, V., et al. (2016b). SNP-Seek II: A resource for allele mining and analysis of big genomic data in Oryza sativa. Current Plant Biology, 7:16–25.

Mather, K. A., Caicedo, A. L., Polato, N., Olsen, K. M., McCouch, S., and Purugganan, M. D. (2007). The extent of linkage disequilibrium in rice (oryza sativa l.). Genetics.

Matsuo, T. (1952). Genecological studies on cultivated rice. Bull. Natl. Inst. Agr. Sci. Jpn. D, 3:1–111.

McCouch, S. R., Wright, M. H., Tung, C.-W., Maron, L. G., McNally, K. L., Fitzgerald, M., Singh, N., DeClerck, G., Agosto-Perez, F., Korniliev, P., et al. (2016). Open access resources for genome-wide association mapping in rice. Nature communications, 7:10532.

Mizuta, Y., Harushima, Y., and Kurata, N. (2010). Rice pollen hybrid incompatibility caused by reciprocal gene loss of duplicated genes. Proceedings of the National Academy of Sciences, 107(47):20417–20422.

Morinaga, T. (1954). Classification of rice varieties on the basis of affinity. Jpn. J. Breed., 4:1–14.

Morishima, H. and Oka, H.-I. (1981). Phylogenetic differentiation of cultivated rice, xxii. numerical evaluation of the indica-japonica differentiation. Japanese Journal of Breeding, 31(4):402–413.

Naranjo, L., Talón, M., and Domingo, C. (2014). Diversity of floral regulatory genes of japonica rice cultivated at northern latitudes. BMC genomics, 15(1):101.

Oka, H. et al. (1991). Genetic diversity of wild and cultivated rice. Rice biotechnology, pages 55–81.

Price, A. L., Helgason, A., Thorleifsson, G., McCarroll, S. A., Kong, A., and Stefansson, K. (2011). Single-tissue and cross-tissue heritability of gene expression via identity-by-descent in related or unrelated individuals. PLoS genetics, 7(2):e1001317.

Pritchard, J. K., Stephens, M., and Donnelly, P. (2000). Inference of population structure using multilocus genotype data. Genetics, 155(2):945–959.

Schatz, M. C., Maron, L. G., Stein, J. C., Wences, A. H., Gurtowski, J., Biggers, E., Lee, H., Kramer, M., Antoniou, E., Ghiban, E., et al. (2014). Whole genome de novo assemblies of three divergent strains of rice, Oryza sativa, document novel gene space of aus and indica. Genome biology, 15(11):506.

Scheipl, F., Greven, S., and Kuechenhoff, H. (2008). Size and power of tests for a zero random effect variance or polynomial regression in additive and linear mixed models. Computational Statistics Data Analysis, 52(7):3283–3299.

Song, Y. H., Shim, J. S., Kinmonth-Schultz, H. A., and Imaizumi, T. (2015). Photoperiodic flowering: time measurement mechanisms in leaves. Annual review of plant biology, 66:441–464.

Stein, J. C., Yu, Y., Copetti, D., Zwickl, D. J., Zhang, L., Zhang, C., Chougule, K., Gao, D., Iwata, A., Goicoechea, J. L., et al. (2018). Genomes of 13 domesticated and wild rice relatives highlight genetic conservation, turnover and innovation across the genus Oryza. Nature genetics, 50(2):285.

Takano, M., Inagaki, N., Xie, X., Yuzurihara, N., Hihara, F., Ishizuka, T., Yano, M., Nishimura, M., Miyao, A., Hirochika, H., et al. (2005). Distinct and cooperative functions of phytochromes a, b, and c in the control of deetiolation and flowering in rice. The Plant Cell, 17(12):3311–3325.

Terao, H. and Mizushima, U. (1942). Some considerations on the classification of Oryza sativa L. into two subspecies, so called Japonica and Indica. Jpn. J. Bot., 10:213–258.

Trapnell, C., Pachter, L., and Salzberg, S. L. (2009). Tophat: discovering splice junctions with rna-seq. Bioinformatics, 25(9):1105–1111.

VanRaden, P. M. (2008). Efficient methods to compute genomic predictions. Journal of dairy science, 91(11):4414–4423.

Walia, H., Wilson, C., Zeng, L., Ismail, A. M., Condamine, P., and Close, T. J. (2007). Genome-wide transcriptional analysis of salinity stressed japonica and indica rice genotypes during panicle initiation stage. Plant molecular biology, 63(5):609–623.

Wang, W., Mauleon, R., Hu, Z., Chebotarov, D., Tai, S., Wu, Z., Li, M., Zheng, T., Fuentes, R. R., Zhang, F., et al. (2018). Genomic variation in 3,010 diverse accessions of asian cultivated rice. Nature, 557(7703):43.

Wang, X., Kudrna, D. A., Pan, Y., Wang, H., Liu, L., Lin, H., Zhang, J., Song, X., Goicoechea, J. L., Wing, R. A., et al. (2014). Global genomic diversity of Oryza sativa varieties revealed by comparative physical mapping. Genetics, pages genetics–113.

Yoshida, S., Forno, D., Cock, J., and Gomez, K. (1976). Laboratory manual for physiological studies of rice, 3rd edn manila: International rice research institute.

Yu, J., Hu, S., Wang, J., Wong, G. K.-S., Li, S., Liu, B., Deng, Y., Dai, L., Zhou, Y., Zhang, X., et al. (2002). A draft sequence of the rice genome (Oryza sativa L. ssp. indica). Science, 296(5565):79–92.

Yuan, J., Chen, D., Ren, Y., Zhang, X., and Zhao, J. (2008). Characteristic and expression analysis of a metallothionein gene, osmt2b, down-regulated by cytokinin suggests functions in root development and seed embryo germination of rice. Plant Physiology, 146(4):1637–1650.

Zhao, K., Tung, C.-W., Eizenga, G. C., Wright, M. H., Ali, M. L., Price, A. H., Norton, G. J., Islam, M. R., Reynolds, A., Mezey, J., et al. (2011). Genome-wide association mapping reveals a rich genetic architecture of complex traits in Oryza sativa. Nature communications, 2:467.

